# Improved Kinect sensor based motion capturing system for gait assessment

**DOI:** 10.1101/098863

**Authors:** Björn Müller, Winfried Ilg, Martin A. Giese, Nicolas Ludolph

## Abstract

Optical motion capturing systems are expensive and require substantial dedicated space to be set up. On the other hand, they provide unsurpassed accuracy and reliability. In many situations however flexibility is required and the motion capturing system can only temporarily be placed. The Microsoft Kinect v2 sensor is comparatively cheap and with respect to gait analysis promising results have been published. We here present a motion capturing system that is easy to set up, flexible with respect to the sensor locations and delivers high accuracy in gait parameters comparable to a gold standard motion capturing system (VICON). Further, we demonstrate that sensor setups which track the person only from one-side are less accurate and should be replaced by two-sided setups. With respect to commonly analyzed gait parameters, especially step width, our system shows higher agreement with the VICON system than previous reports.

## Introduction

The Microsoft Kinect v2 sensor is a low-priced depth camera which was originally meant to be used for gaming in combination with the Microsoft Xbox One console. Recently, increasing interest in using the Kinect sensor for general purpose motion capturing (MoCap) of humans has emerged, especially for clinical and scientific motion analysis of gait [1–6]. Due to the low costs it was proposed to utilize the Kinect sensor as a cost-efficient alternative to expensive gold standard motion capturing systems [1, 7]. A similar attempt has been made using the first generation of the Kinect sensors (Kinect for Xbox 360) which was designed to be used with the Microsoft Xbox 360 console [3, 8–10]. However, the Kinect for Xbox 360 relies on the recognition of reflected infrared patterns to acquire the depth information and great effort has been put into studying and reducing the interference of the patterns when using multiple sensors [11, 12]. In contrast to this, the Kinect v2 uses time of flight measurements, is less sensitive to interference with other sensors and provides a higher resolution. The term “time of flight” describes the method to determine the distance to an object by measuring the time a laser pulse needs to travel from the sensor to the object and back. The Kinect v2 sensor has a horizontal field of view of about 70 degrees and can cover 4.5 meters in depth reliably. Due to the limited size of the tracking volume of the Kinect sensor, single sensor approaches were mostly constrained to examinations of body posture and balance during stance or of walking on a treadmill [2, 5, 6]. In order to cover a larger volume, setups with multiple Kinect sensors have been proposed [1, 3, 7, 13].

With the Kinect v2 software development kit (SDK) Microsoft provides an easy way to access the different data streams of the sensor. The most important data streams for the purpose of motion tracking are the color, depth and skeleton streams. In a previous study it has been described how these streams can be utilized to spatially calibrate multiple sensors [14]. A more clinically motivated study examined successfully 10-meter walking using four Kinect v2 sensors. The sensors were lined up on the left side of the walking corridor. Based on the averaged joint position estimates, several gait parameters have been extracted and compared to a gold standard MoCap system [1]. The depth resolution of the Kinect v2 sensor, however, depends not only on the distance but also on the view angle from which a plane is measured [15]. In addition, the error of the joint position estimation algorithm increases with the view angle which is likely caused by partial self-occlusion [4]. Motion capturing from only one side using Kinect sensors, might therefore introduce biases and unnecessary inaccuracies in the estimation of joint positions.

The aim of this study was to (1) develop a scalable motion tracking system based on Kinect v2 sensors, (2) to examine in how far one-sided tracking biases gait parameters and (3) to propose a camera setup which circumvents the potential drawbacks of one-sided tracking. In order to evaluate the quality of our system, we conducted a statistical comparison of the tracking performance with a VICON MoCap system based on the gait parameters: step length, step width, step time, stride length and walking speed. Six Kinect v2 sensors were used to cover a walking corridor of more than six meters. We put emphasis on a detailed description of the system since even though several Kinect-based MoCap systems have been described, no standard has been defined yet.

## Methods

### Kinect Sensor

Microsoft’s Kinect v2 provides five video related data streams [16]. Besides the color (1920x1080@30Hz) and infrared (512×424@30Hz) data streams, the Kinect provides depth images (512×424@30Hz), body index images (512x424@30Hz) and the skeleton information for every tracked person (25joints@30Hz). The sensors tracking volume is defined by the field of view (FOV, 70°horizontally, 60° vertically) and the range of depth sensing (0.5–4.5 meters).

These data streams can be accessed using Microsoft’s software development kit (v2.0). Color images are provided with 4 bytes per pixel (BGRA) and depth images with 2 bytes per pixel resolution. In order to distinguish tracked persons, the Kinect SDK assigns indices which are stored in body index images and take one byte per pixel. The joint positions are provided at a resolution of 4 bytes per coordinate (12 bytes per joint). Every frame contains a timestamp representing the local time of the computer. Besides the transformation between the pixel coordinate systems of the data streams (e.g., the color and depth data streams have different resolutions), the SDK can also be used to translate depth images into 3d point clouds (inverse perspective projection). This way it is possible to acquire the color values for every depth image pixel and display a colored 3d point cloud (Figure 2 A, B).

According the Microsoft’s specifications each Kinect v2 sensor requires a dedicated USB 3.0 controller. Additionally, even though Microsoft initially planned to support multiple Kinect sensors per computer, the current SDK version (v2.0) does not support this feature. Thus, each Kinect sensor has to be connected to a dedicated computer.

### Hardware & Software Architecture

Our hardware architecture consists of six Microsoft Kinect v2 sensors which are each plugged into a separate mini-computer (Zotac Zbox ID83 Plus, Intel Core i3 dual core 2.5GHz, 8GB Ram, 256GB SATA-3 SSD, Windows 10 Pro) and a dedicated computer for control and monitoring (Figure 1). All computers are connected via a gigabit Ethernet network. In order to reduce network traffic while recording, we decided to store the data locally instead of transmitting the data directly to a remote computer. To this end, solid state drives have been used. These provide a higher write speed than conventional hard disk drives. We mounted the Kinect sensors on and strapped the mini-computers to tripods for a solid stance and flexible setup. As dedicated computer for controlling and monitoring the system, we used a laptop computer with a dedicated 3d graphics card for visualization.

**Figure 1:**
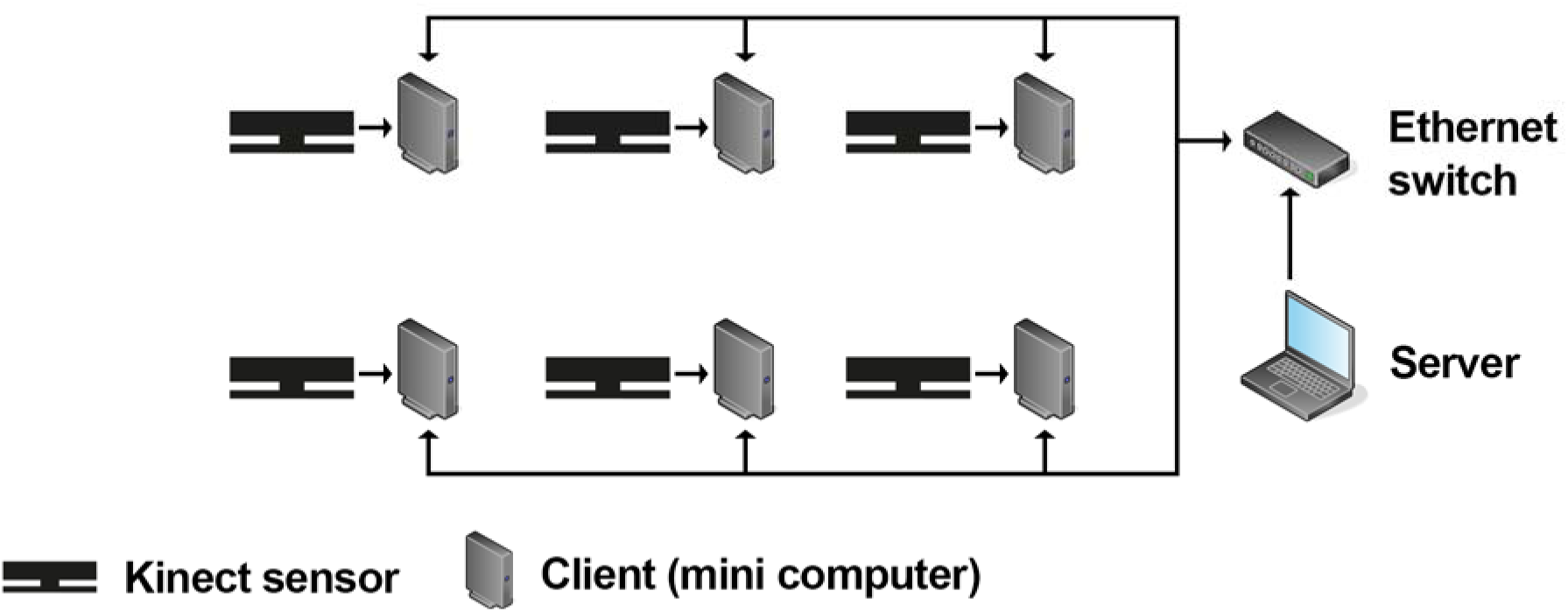
Connections between the Kinect sensors, Zotac mini computers running the client software and the server computer in a setup with six Kinect sensors.

Based on this hardware architecture we implemented a client server software architecture (Figure 1) such that each mini-computer (in the following called: client) runs the client software and the desktop computer (in the following called: server) runs the server software. Both components were implemented using C# in Microsoft’s .NET framework (v4.5). Communication between the server and its clients is based on the TCP/IP protocol and a custom-made software-level protocol which defines the format for the transfer of commands and recorded data.

The Kinect data streams recorded by the clients had to be synchronized in time. To this end we initially synchronized the clocks of all clients with the server using Microsoft Windows’ time service. However, we noticed that, even when the computers’ clocks had been synchronized in this way, they differed by several hundreds of milliseconds and sometimes even by seconds. In order to synchronize the computer clocks more precisely, we used Greyware’s DomainTime II (Greyware Automation Products, Inc.) which implements the precision time protocol (PTP). PTP was designed for time critical applications, e.g. in industry, and allows to synchronize computers in a network with millisecond accuracy. We were thereby able to achieve a maximum difference of two milliseconds between the clocks of all involved computers. This difference has been monitored before and during every recording using Greyware’s monitoring software in order to ensure synchrony. The timestamps of the captured frames have then been used to align the data streams in time.

### Server & client software

The server software consists of several modules which are responsible for the data management, recording, live-view and spatial calibration of the system. The data management module structures the data hierarchically in projects, subjects, sessions and recordings. Additionally, it keeps track of the recorded data and its location. Since each client stores the data locally and sends it only on request to the server via Ethernet, the data management module also identifies and prevents data inconsistencies, such as incomplete or partial transmission.

The recording module realizes the synchronous start and stop of recording for all clients. Due to network transmission the clients might receive the commands at slightly different times. To counteract the resulting problem, first of all, each client is buffering the two most recent seconds of all data streams and secondly, when a recording is started, the transmitted command contains the server’s current timestamp. Thereby, even when the clients receive the start command at a slightly different time, they can compensate for this using the timestamp and buffered data. During recording the data is stored locally by each client using custom binary data formats. The stop procedure is implemented in a similar way in order to make sure that all clients record for the same duration.

Within the live-view module, the overall 3d scene (merged point clouds of all clients, Figure 2 A, B) and the depth images (Figure 2 C) can be displayed. While the on-demand visualization of the overall 3d scene serves mere the purpose of illustration, displaying the depth images is helpful to facilitate the process of setting up the system. Tracked bodies are highlighted such that one can easily identify blind-spots in the tracking volume. Thereby, blind-spots and too little overlap of the sensors’ tracking volumes can be easily avoided when positioning and orienting the sensors. For the 3d visualization, every client transforms the depth images into 3d point clouds in real time using the Kinect SDK. The current point cloud can be requested by the server for visualization and spatial calibration (see Spatial calibration). In order to perform a reconstruction of the whole scene, the server requests the point cloud from every client, transforms these into the global coordinate system using the spatial calibration parameters (see Spatial calibration) and visualizes every point as small cube in an OpenGL viewport (Figure 2 A). Each cube is colored according to the corresponding pixel in the color image.

**Figure 2:**
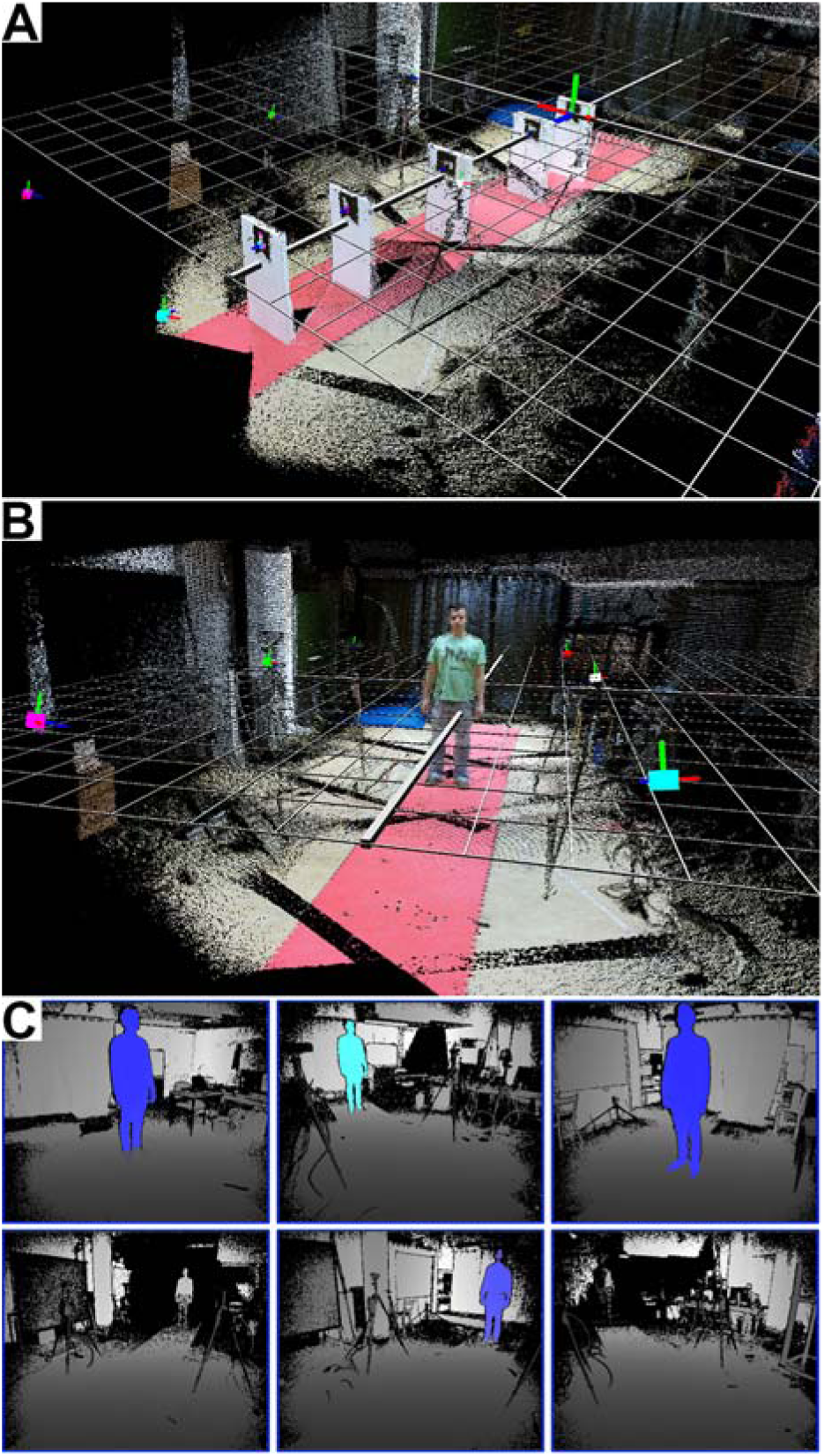
(A-B) Three-dimensional point clouds of six Kinect sensors after spatial calibration. Each point is represented as tiny cube with the color of the corresponding the pixel in the color image. The big colored cubes indicate the Kinect sensors. The red, green and blue lines attached to the colored boxes (sensors) indicate the axes of local coordinate system. Notice that the edge of the red coating on the floor is very straight, which illustrates the precise spatial calibration. (A) The marker in the very back is the marker with id 1 and represents the origin of the global coordinate system which is indicated by the grid. (B) A person standing in the tracking volume to visualize the dimensions. (C) Screenshot of the live depth images view. The tracked body is highlighted. By walking through the tracking volume one can easily identify blind spots.

### Spatial calibration

The spatial calibration is equivalent to estimating the position and orientation of every Kinect sensor in the global coordinate system. Inspired by the work of Kowalski et al. [14], we use two-dimensional markers which can easily be detected in the color images captured by the Kinect sensor (see Figure 3 A). Using these markers, we defined the global coordinate system. In some setups however, the sensors might be so far away from each other, that not every sensor sees all markers. We extended their solution by a flexible concatenation of Euclidian transformations (e.g., rotations and translations) in order to overcome this problem. Thus, not every marker needs to be visible to every sensor and the spatial relation between the markers does not need to be known prior to calibration. The calibration procedure consists of six steps which are described in more detail in the following.

**Figure 3:**
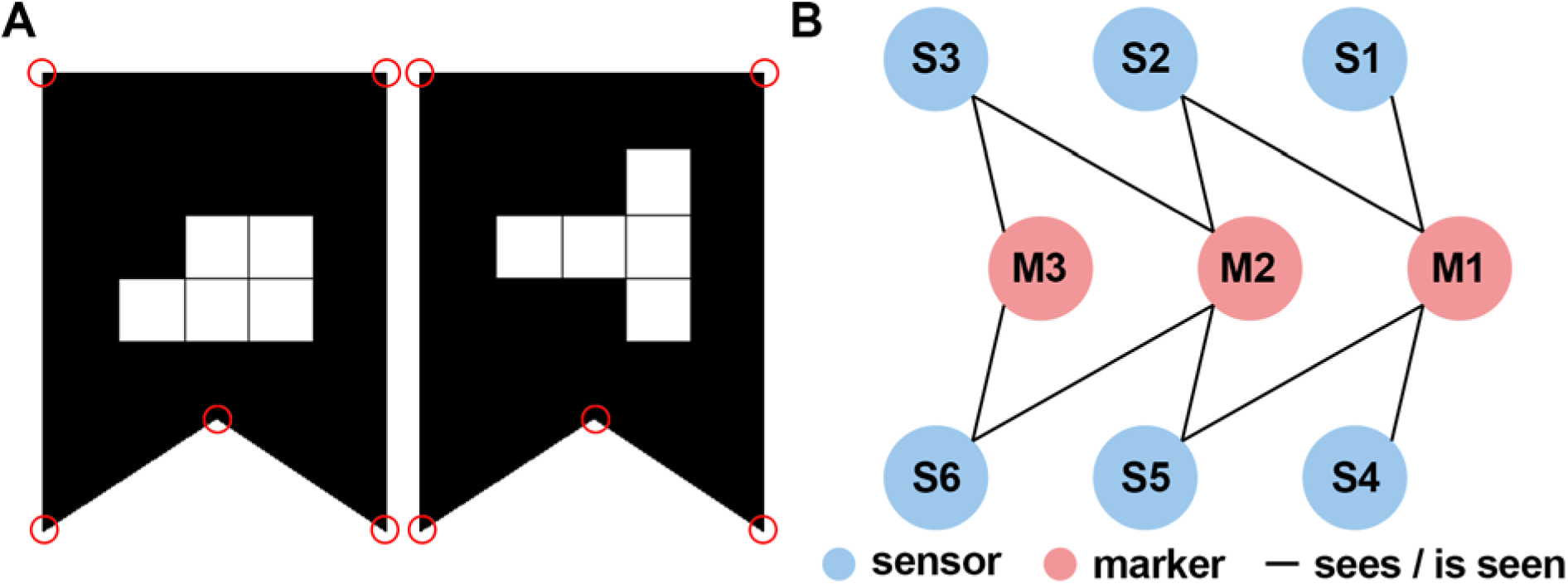
(A) Makers used for the spatial calibration of the system. The shape is easily detectable in the RGB image; the white squares in the center encode the marker id [14]. The red circles indicate the salient points which have been used for defining the position and orientation of the marker. (B) Graph illustrating the “sees / is seen” relation (edges) between sensors (blue vertices) and markers (red vertices). For example, S3 cannot see M1 directly but indirectly via M2 and S2. A second possibility is via M2 and S5.

The two-dimensional markers proposed by Kowalski et al. [14] consists of a salient shape (rectangle with a dent at the bottom) and a unique pattern of white squares in a 3x3 grid (Figure 3 A). In the first step, we find these shapes in the color image using the OpenTK library and then interpret the pattern as identification code (for details see Kowalski et al. [14]). Besides identifying the markers uniquely, the identification code also breaks the symmetry of the shape. In the second step, we use the point cloud to determine the 3d coordinates of the salient points (corners) belonging to the marker. While the third step consists only of the transmission of the information about the markers to the server, we apply a Procrustes analysis [17] to the 3d positions of the salient marker points in the fourth step. The result of the Procrustes analyses are estimates of the relative positions and orientations (Euclidian transformations) between the markers and sensors.

The fifth step is to determine the position and orientation of every sensor relative to the marker with id 1. This marker can be placed anywhere in the tracking volume and defines the origin and directions of the axes. Since the Kinect sensor has a limited field of view and can only estimate depth values of about 4.5 meters accurately, depending on the setup, the marker is not in the field of view of all sensors. Notice, that in the fourth step only the relative position and orientation between markers and sensors that are visible to each other have been determined. In order to tackle this problem, we developed an algorithm which is flexible with respect to the placement of the calibration markers. Our algorithm is based on the idea, that even if the marker that represents the origin is not visible to a certain sensor, the relative position and orientation could still be calculated as concatenation of the Euclidian transformations which express the relative positions and orientations of the other sensors and markers (Figure 3 B). For example, if sensor 1 sees the markers 1 and 2 but sensor 2 sees only marker 2, the position and orientation of sensor 2 relative to marker 1 can be calculated using the relative position and orientation between the two sensors which can be determined using marker 2. In other words, marker 1 is indirectly visible to sensor 2. In order to find this concatenation of Euclidian transformations automatically, we use a bipartite undirected graph *G* = (*V,E*) with *V* = *M* ∪ *S*, *M* ∩ *S* = ∅ and *E* = {(*s, m*) ∈ *S* × *M*, (*m, s*)∈ *M* × *S* | “sensor s sees marker m”} where *M* denotes the set of all markers and *S* the set of all sensors (Figure 3 B). Thus, we can verify that every sensor can see the origin (in-)directly by evaluating whether the graph is connected. Furthermore, the position and orientation of every sensor relative to the origin can be determined by finding a path between the two respective nodes in the graph (see Supplementary Material for details), because every path in the graph describes a concatenation of Euclidian transformations. In the case that there are multiple paths, we use the shortest one, because every transformation is based on an estimation that includes an estimation error and thus reduces the accuracy of the final estimate.

Having hereby determined a coarse estimate of the sensor positions and orientations relative to the origin (see Supplementary Material for details), we perform a refinement using the iterative closest point method in the sixth step (see [14] for details). Using these refined estimates, we can merge the 3d scenes (Figure 2 A, B) and Kinect skeletons in the global coordinate system.

### Evaluation of the gait analysis system

In order to evaluate the quality of the developed MoCap system we compared gait parameters measured by our system to measurements gathered using a VICON system (Vicon Motion Systems Ltd). VICON motion capturing systems are seen as gold standard in optical gait analysis [18]. We operated both systems in parallel, allowing us to compare the two systems based on individual steps.

### Subjects and task

We recorded 10 healthy subjects performing a 7-meters walk at comfortable speed ten times. Subjects were wearing tight clothes and normal shoes without heels. Subjects’ age ranged from 18 to 35 years. All subjects gave written informed consent prior to participation. The experimental procedure was approved by the local ethical review board of the University Clinic in Tuebingen.

### VICON motion capturing system

As ground truth for the accuracy evaluation of our system we used a VICON MX motion capture system with 10 cameras. The VICON system is an optical tracking system which tracks three-dimensional movement trajectories of reflective markers with up to 1mm accuracy. We used this system to track 12 markers attached to the hip, legs and feet according to VICON’s Plugin-Gait marker-set for lower body measurements at a temporal resolution of 120 Hz. Our VICON setup is able to track about 6 meters in length with high precision. Marker trajectories were recorded and processed using the commercial software Nexus (v2.2, Vicon Motion Systems Ltd).

### Kinect sensor setup for gait analysis

Six Kinect sensors were arranged as an avenue (Figure 4) in order to cover the tracking volume of the VICON system. We placed the sensors pairwise in rows along the walking direction for two reasons: (1) reduction of self-occlusions of the tracked person and (2) more accurate tracking. The theoretical length of the tracking volume is about 9 meters of which we used the overlapping 6 meters with the VICON setup.

**Figure 4:**
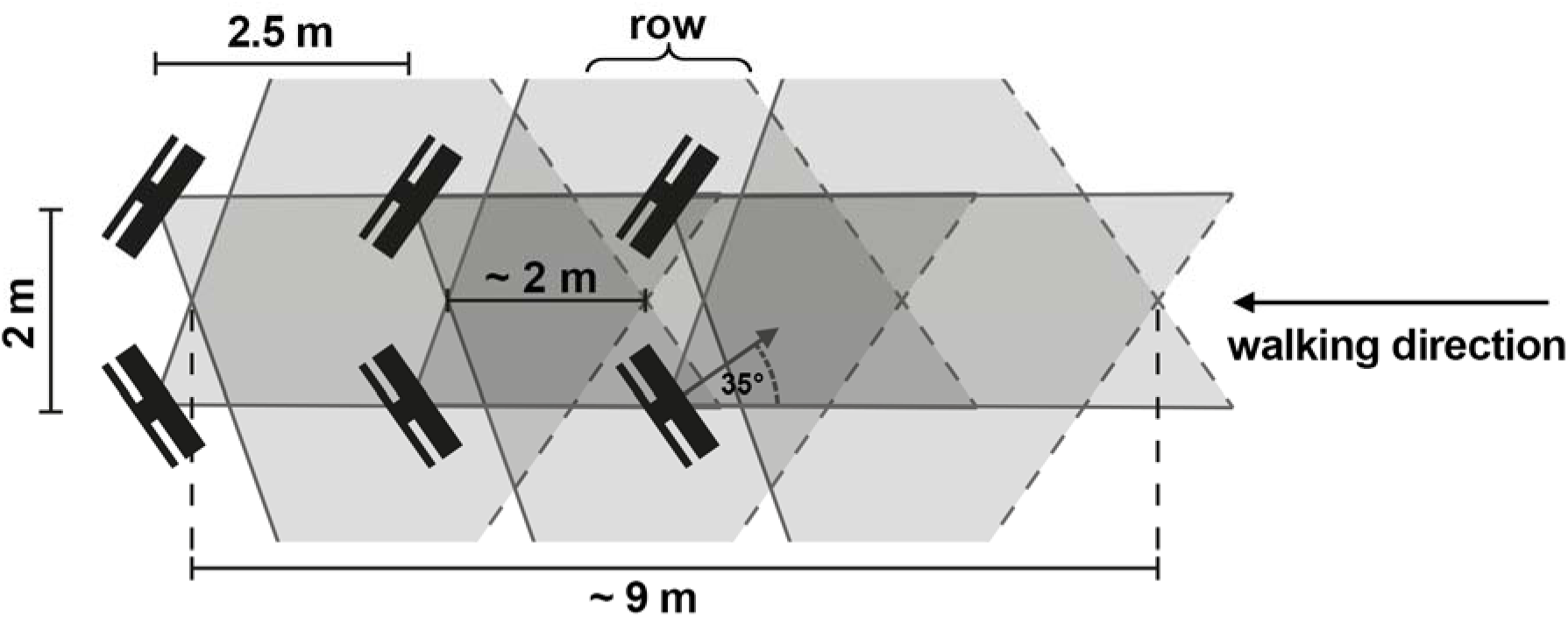
Arrangement of the Kinect sensors illustrating the overlapping tracking volumes. Sensors were arranged using this pattern but then spatially calibrated to achieve high precision. The theoretical length of the tracking volume in walking direction using this setup is about 9 meters. The Kinect sensors were arranged in a way ensuring that the VICON tracking volume was completely covered.

We noticed during initial tests of our system that the Kinect skeleton fitting algorithm depends heavily on the view angle from which the sensor tracks the body (Figure 5 A, B, see Supplementary Material). A similar finding has previously been reported [4], however the cause of the inaccuracy had not been examined in detail before. Tracking a person only from one side, e.g. from the left side, could degrade the tracking precision and potentially bias the skeleton fitting. Additionally, when tracking a walking person only from one side, the leg of the opposite body half is periodically occluded during the gait cycle. In these situations, the Kinect skeleton fitting algorithm initially tries to infer the position of the occluded joints and if that fails the respective joints are labeled as untracked. For the usual application of the Kinect sensor in gaming this is not an issue, since the accuracy is less important. However, for gait analysis, we do not want to rely on the inferred joint positions or biased skeleton fits. Therefore, we ignore joint positions which are labeled as untracked and record the person from two sides which allows us to correct biases and to reduce self-occlusions.

**Figure 5:**
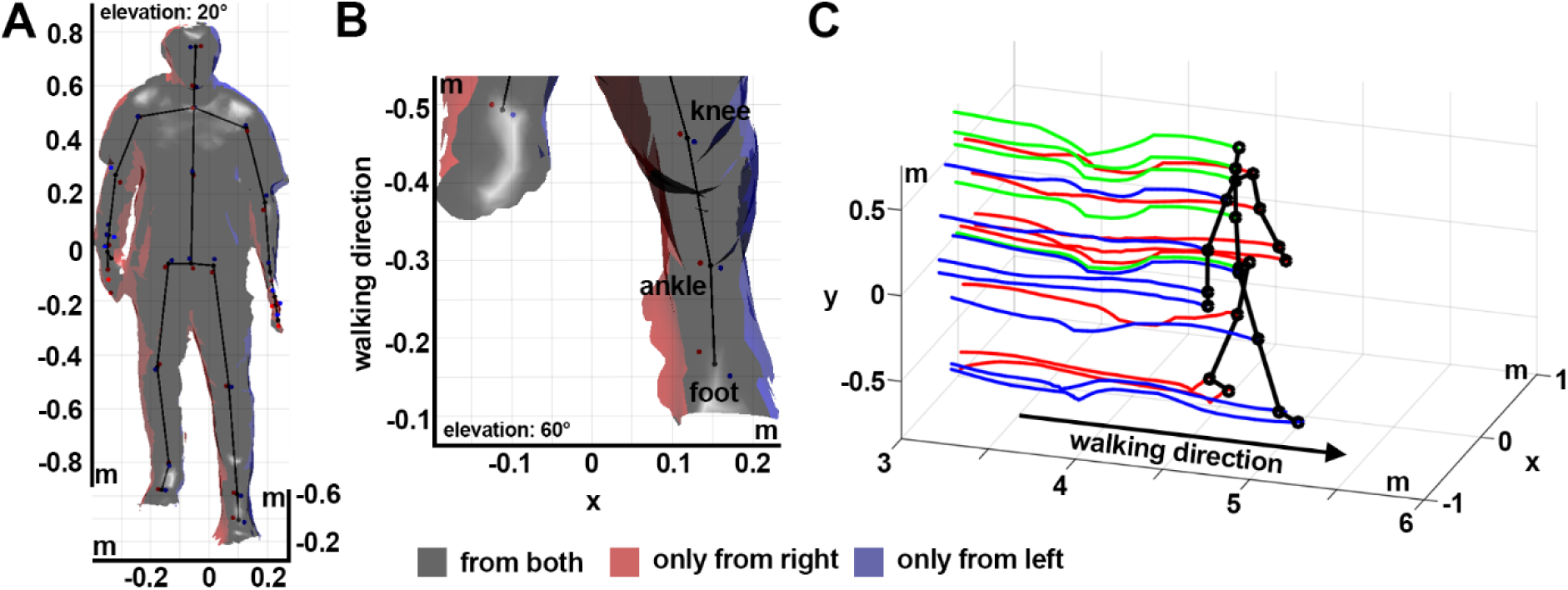
(A, B) Three-dimensional reconstruction of the body surface and the corresponding skeleton reconstruction using two sensors. The surface was estimated based on the 3d point clouds using the marching cubes algorithm in MeshLab [22]. Surface areas tracked by only one of the two sensors are highlighted in red (right) and blue (left). Corresponding Kinect skeleton joint positions estimates of the two sensors are shown as red and blue dots. The spatially averaged skeleton is indicated as black stick figure. (B) Magnification of the left lower leg. Notice, that the joint positions estimates of the left sensor (blue) are closer to the surface which is only tracked by the left sensor (blue), correspondingly for the joint positions estimates of the right sensor. (C) Averaged skeleton and joint position trajectories during walking obtained using six sensors.

We also made sure that the tracking volumes of two rows overlap by about two meters along the walking path (Figure 4). The Kinect pose estimation algorithm needs some time to recognize a person that enters the volume. Hence, we had to make sure that the sensors in the next row are already tracking the person when s/he is about to leave the tracking volume of the previous sensor pair. We empirically estimated that an overlap of the tracking volumes of about two meters is sufficient for normal walking speed. Thus, the sensor pairs were placed 2.5 meters apart along the walking path. Within each row the sensors were rotated inwards by about 35 degrees and placed two meters apart which provides plenty of width for walking in the corridor (Figure 4).

## Data analysis

Before being able to compare the performance of both tracking systems we had to pre-process the data. First, we had to integrate the skeleton information of the different Kinect sensors. Secondly, we had to extract the gait-features from both data sets (VICON and Kinect).

### Integrating the information gathered by different Kinect sensors

The information gathered from the different Kinect sensors was integrated in space and time. Integration in space was performed using the Euclidian transformations that were computed as result of the spatial calibration. The clocks of all involved computers were synchronized using the PTP protocol, which guarantees that frames with the same timestamp were captured at approximately the same time. However, the Kinect sensor captures frames at a slightly varying frequency of about 30 Hz. We resampled the recordings with a fixed sampling rate of 30 Hz using linear interpolation. Since we do not have any information about the differences in sensor noise, we weighted all sensors equally and calculated the average across all skeletons tracked by the different sensors for every sample. Untracked or inferred joint positions were considered as missing values. The result of this integration is a spatially averaged skeleton across time. For the subsequent analysis, this procedure was performed once using all Kinect sensors (left and right) and once using only the Kinect sensors tracking the person from the left. Subsequently, we filtered the three-dimensional joint trajectories in time for both systems. The resulting three-dimensional Kinect skeleton trajectories are exemplified in Figure 5 C.

### Analyzed gait features

In order to compare the tracking accuracy of gait, we extracted five parameters from the two datasets (VICON and Kinect) independently: walking speed, step length, stride length, step width and step time. To this end we examined individual steps by first identifying foot placements (Figure 6) in each recording. In order to identify the foot placements we determined the events when one foot passes the other in walking direction (see Figure 7). These events are well-defined and easy to detect by searching for the intersections of the ankle trajectories (black crosses in Figure 7). Subsequently, the stride length was calculated as the distance between subsequent foot placements of the same foot [19]. The step time and length describe the time passed, and the distance between the two feet in walking direction, whereas the step width describes the lateral distance between the feet, at the time of a foot placement (Figure 6). The walking speed was determined by dividing the sum of all step lengths by the sum of all step times within each recording. Since the first step is usually significantly smaller due to the necessary acceleration, we asked subject to begin walking with the left foot and then excluded the first step on the left as well as the first stride on the right side.

**Figure 6:**
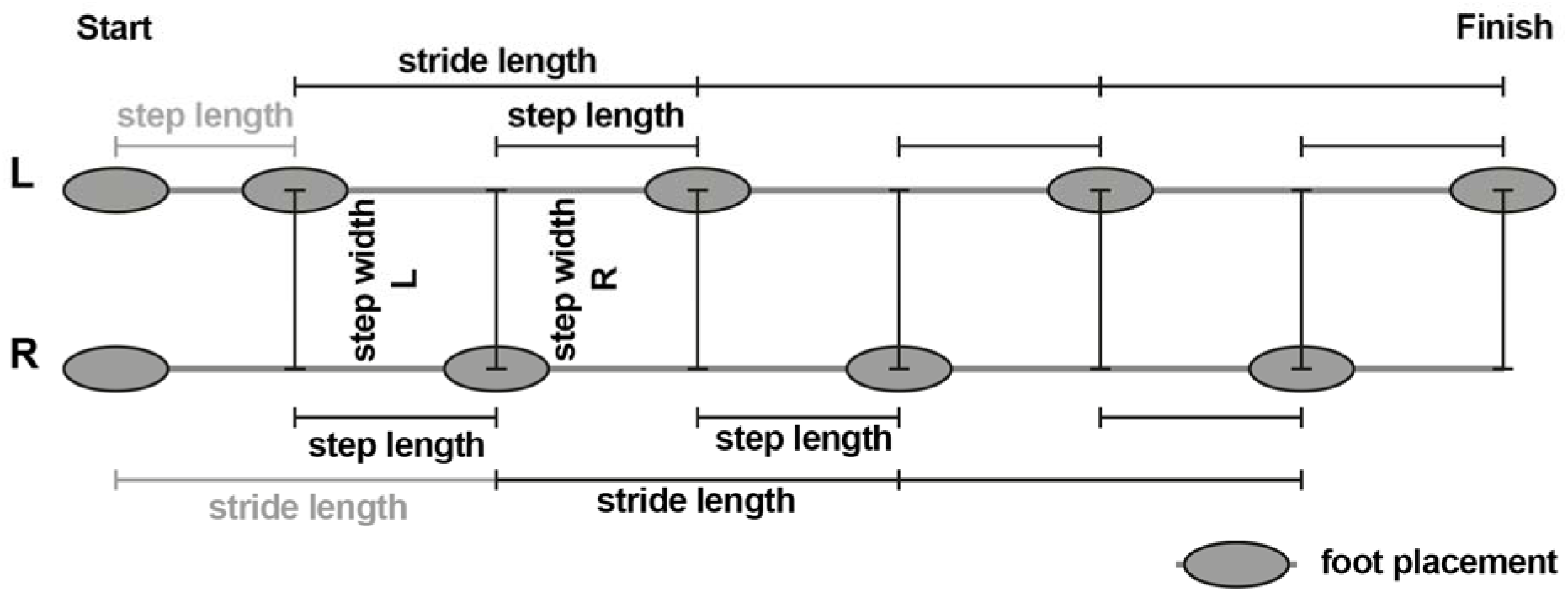
Illustration of the analyzed spatial gait parameters. Subjects were asked to always start walking with their left foot. The first step length left and first stride length right (gray) were excluded from the analysis, since these are generally shorter than the steps during actual walking (acceleration phase). Subjects did not stop at the end of the track volume (finish) so that there was no slowing down. Depending on the subjects’ individual step lengths there might be an unequal number of left and right steps. Parameters are assigned to the left or right foot depending on which foot was last placed on the floor, e.g. the first step length and step width are assigned to the left foot.

**Figure 7:**
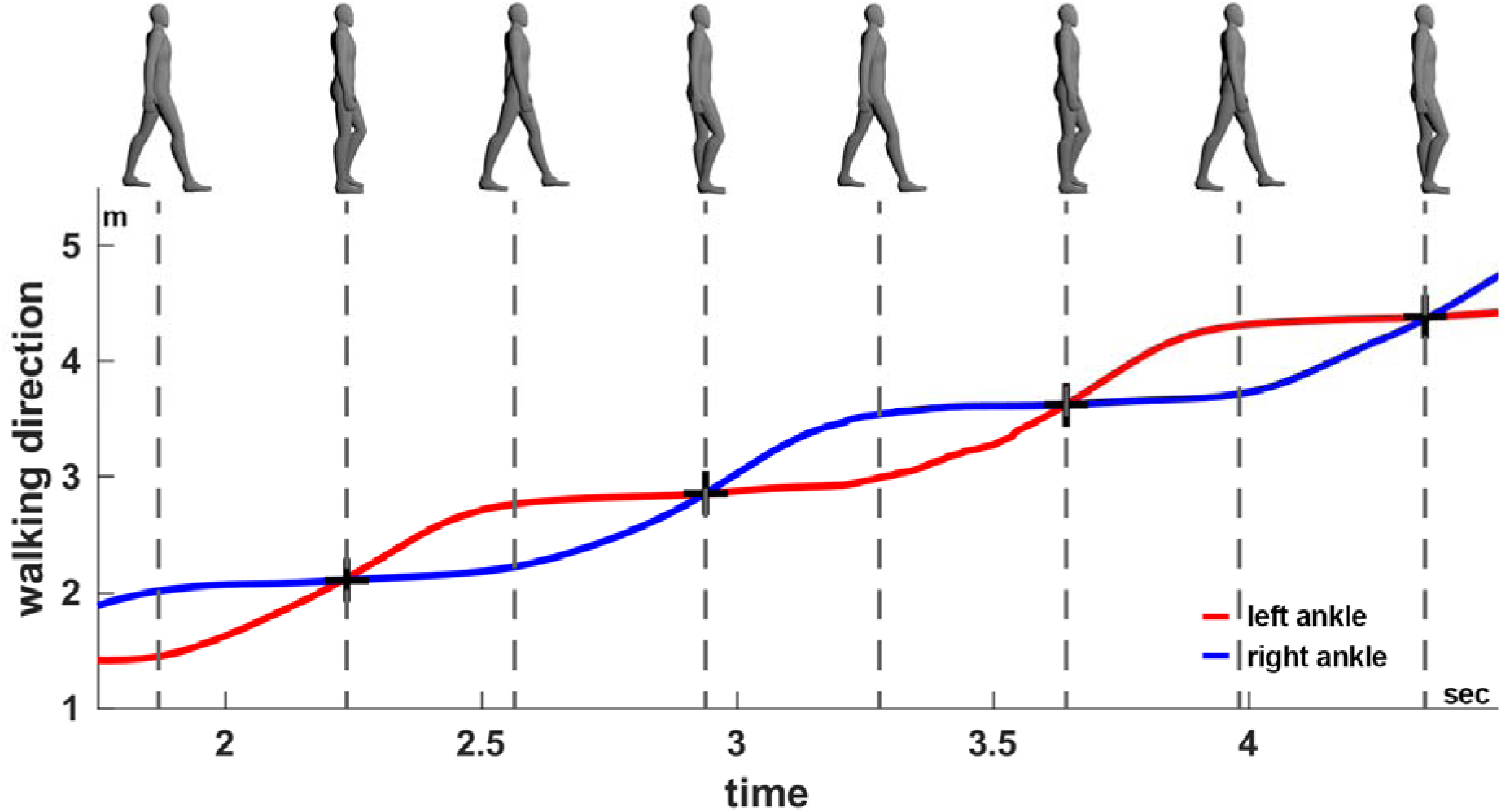
Foot events (black crosses) based on the ankle trajectories, here exemplified using the Kinect skeleton averaged across all sensors. Using the same procedure foot events were extracted from the VICON data. Top: Snapshots of the body posture during walking, bottom: ankle position of the left (red) and right (blue) foot in walking direction.

### Quantitative comparison of the tracking results

The agreement between the Kinect-based gait analysis system and the VICON system has been evaluated using Pearson’s correlation coefficient, Bland-Altman’s method for assessing the agreement between two clinical measurement methods [20] and the intraclass correlation coefficient for absolute agreement (ICC(A,1)) on three levels of detail: (1) single steps/strides left and right, (2) averaged steps/strides left and right per subject and (3) average steps/strides per subject (pooled over left and right steps/strides). Bland-Altman’s method is primarily a graphical analysis.

However, it provides three well-interpretable parameters: (1) bias (average difference between measurement methods), (2) reproducibility coefficient (RPC, standard deviation of the difference between the measurement methods) and (3) coefficient of variation (CV, standard deviation of the difference between the measurement methods divided by the average measurement) in percent. While a non-zero bias indicates a systematic deviation, the RPC represents the overall variability between the two methods. The CV quantifies the variability in terms of the average measurement value and is therefore better suited when comparing measures with different mean values. ICC(A,1) takes values between zero and one, where one is perfect agreement. Following [21], we classified its value according to the categories: poor (0–0.4), fair (0.4–0.59), good (0.6–0.74) and excellent (0.75–1.0) absolute agreement.

Motivated by the observation that the joint positions measured by the Kinect sensors depend on the view angle, we performed two gait analyses for the Kinect system: (1) using only the sensors from the left side in walking direction (one-sided) and (2) using all sensors (two-sided). The resulting gait parameters were separately compared to those measured using the VICON system. Instead of recording every subject twice, we used the same recordings but reconstructed the skeleton using either only the left Kinect sensors or all of them. Since the step time and walking speed do not depend on the view angle, we compared the Kinect and VICON measurements of these parameters without distinguishing between one- and two-sided Kinect tracking.

## Results

### Agreement of temporal gait parameters

The summary and agreement statistics of the view angle independent gait parameters, walking speed and step time, are listed in Table 1. For both measures the intraclass correlation coefficients for absolute agreement ICC(A,1) are excellent on all detail levels. Best agreement has been found for the averaged step time with an ICC(A,1) close to one, zero bias and RPC close to zero. High agreement was also found for the step time on the other detail levels (single step times, subjects’ average step times left/right). Despite excellent agreement for the walking speed according to the intraclass correlation coefficient for absolute agreement, we observe a small bias of 0.53cm/s. Overall, these results indicate excellent agreement between the evaluated Kinect system and VICON in the temporal gait parameters.

**Table 1:** Summary and agreement statistics for the view angle independent gait parameters walking speed and step time. Abbreviations. RPC:reproducibility coefficient (1.96*SD); CV: coefficient of variation (SD of mean values in %); ICC(A,1): intraclass correlation coefficient for absolute agreement. Subject averages across steps (AV) and sides (AVG) are reported for comparison with previous publications. Measures in bold can be compared to previously reported systems [1, 2].

| Parameter | Our System | VICON | Pearson's correlation |  |  | Bland-Altman |  |  | ICC(A,1) |
| --- | --- | --- | --- | --- | --- | --- | --- | --- | --- |
| | Mean ± SD | Mean ± SD | R | P | N | Bias | RPC | CV (%) | |
| **AV Walking Speed (cm/s)** | **123.06 ±14.26** | **122.53 ±14.50** | **1.000** | **<0.001** | **10** | **0.53** | **0.80** | **0.33** | **0.999** |
| Step time L (s) | 0.60 ±0.07 | 0.61 ±0.07 | 0.927 | <0.001 | 267 | -0.00 | 0.05 | 4.49 | 0.924 |
| Step time R (s) | 0.60 ±0.07 | 0.60 ±0.07 | 0.927 | <0.001 | 281 | 0.00 | 0.05 | 4.38 | 0.925 |
| AV step time L (s) | 0.60 ±0.07 | 0.61 ±0.07 | 0.979 | <0.001 | 10 | -0.01 | 0.03 | 2.29 | 0.978 |
| AV step time R (s) | 0.60 ±0.06 | 0.60 ±0.06 | 0.976 | <0.001 | 10 | 0.00 | 0.03 | 2.33 | 0.976 |
| **AV step time AVG (s)** | **0.60 ±0.06** | **0.60 ±0.07** | **1.000** | **<0.001** | **10** | **0.00** | **0.00** | **0.23** | **1.000** |

### Agreement of spatiotemporal gait parameters using one-sided Kinect setup

We have listed the summary and agreement statistics for the comparison of the one-sided Kinect tracking and VICON in Table 2. For the averages across subjects and feet in step length, stride length, and step width we find excellent agreements. On the single steps level, we found that the agreement for the right step length and width is worse than for the left side, suggesting overall less precise measurements of the right body half. The worst agreement is found for the step width on the right (step width R) with an ICC(A,1) of 0.297, rather large bias of 2.68 cm and a CV of about 50%. Similarly, the subjects’ average step width on the right (AV step width R) shows only fair agreement ICC(A,1) = 0.452 with a similarly large bias as for the single steps (2.66 cm). Overall, the analysis shows that the agreement for the right body half is worse than for the left, when tracking only from the left side. Even though the ICC for the averaged step width across feet indicates excellent agreement, the agreement is not as good as for the step and stride length, indicating that the inference of the step width is less precise.

**Table 2:** Summary and agreement statistics for spatiotemporal gait parameters using Kinect tracking from the left side (one-sided). Notice, the difference in agreement between the left and right body half for the step width and step length. Abbreviations: RPC: reproducibility coefficient (1.96*SD); CV: coefficient of variation (SD in % of the mean value); ICC(A,1): intraclass correlation coefficient for absolute agreement. Subject averages across steps (AV) and sides (AVG) are reportedfor comparison with previous publications. Measures in bold can be compared to previously reported systems [1, 2].

| Parameter | Our System | VICON | Pearson's correlation |  |  | Bland-Altman |  |  | ICC(A,1) |
| --- | --- | --- | --- | --- | --- | --- | --- | --- | --- |
| | Mean ± SD | Mean ± SD | R | P | N | Bias | RPC | CV (%) | |
| Stride length L (cm) | 145.95 ±11.08 | 145.36 ±8.49 | 0.786 | <0.001 | 267 | 0.59 | 13.43 | 4.71 | 0.758 |
| Stride length R (cm) | 146.27 ±10.22 | 147.04 ±9.01 | 0.923 | <0.001 | 187 | -0.78 | 7.77 | 2.70 | 0.913 |
| Step length L (cm) | 72.85 ±6.18 | 73.95 ±5.36 | 0.667 | <0.001 | 267 | -1.10 | 9.34 | 6.49 | 0.649 |
| Step length R (cm) | 73.40 ±8.36 | 71.67 ±4.48 | 0.538 | <0.001 | 281 | 1.73 | 13.81 | 9.72 | 0.434 |
| Step width L (cm) | 10.02 ±4.69 | 11.51 ±3.15 | 0.569 | <0.001 | 267 | -1.49 | 7.62 | 36.13 | 0.493 |
| Step width R (cm) | 14.58 ±7.33 | 11.91 ±2.97 | 0.475 | <0.001 | 281 | 2.68 | 12.69 | 48.88 | 0.297 |
| AV stride length L (cm) | 146.30 ±8.70 | 145.66 ±7.98 | 0.997 | <0.001 | 10 | 0.64 | 1.90 | 4.71 | 0.991 |
| AV stride length R (cm) | 146.16 ±8.94 | 146.95 ±8.43 | 0.994 | <0.001 | 10 | -0.79 | 2.15 | 2.70 | 0.989 |
| **AV stride length AVG (cm)** | **146.24 ±8.75** | **146.20 ±8.17** | **0.998** | **<0.001** | **10** | **0.04** | **1.53** | **0.53** | **0.996** |
| AV step length L (cm) | 72.97 ±4.17 | 74.10 ±5.13 | 0.939 | <0.001 | 10 | -1.13 | 3.69 | 2.56 | 0.900 |
| AV step length R (cm) | 73.36 ±5.12 | 71.67 ±3.60 | 0.898 | <0.001 | 10 | 1.69 | 4.84 | 3.40 | 0.799 |
| **AV step length AVG (cm)** | **73.19 ±4.40** | **72.82 ±3.96** | **0.996** | **<0.001** | **10** | **0.36** | **1.12** | **0.78** | **0.988** |
| AV step width L (cm) | 10.03 ±2.67 | 11.55 ±1.85 | 0.750 | 0.01 | 10 | -1.52 | 3.47 | 16.39 | 0.591 |
| AV step width R (cm) | 14.57 ±2.44 | 11.91 ±1.74 | 0.845 | 0.002 | 10 | 2.66 | 2.63 | 10.13 | 0.452 |
| **AV step width AVG (cm)** | **12.39 ±2.15** | **11.73 ±1.76** | **0.944** | **<0.001** | **10** | **0.66** | **1.49** | **6.30** | **0.882** |

### Agreement of spatiotemporal gait parameters using two-sided Kinect setup

The statistical comparison of the two-sided Kinect tracking with the VICON results is shown in Table 3. Overall the agreement is much better than for one-sided Kinect tracking. All but one measure (AV step width R) reach good agreement and most show excellent agreement. On the level of single steps, specifically the agreement in step width is much higher than for one-sided tracking (ICC(A,1) is about 0.66 for both sides instead of 0.49 for the left and 0.29 right side). Additionally, the average step width (AV Step width AVG) is much less biased (0.39 cm vs. 0.66 cm) and shows an agreement more similar to the other measures (ICC(A,1) = 0.910). These results demonstrate the gain in precision due to twosided tracking.

**Table 3:** Summary and agreement statistics for spatiotemporal gait parameters using Kinect tracking from the left and right side (two-sided). Notice in comparison to Table 2, that the difference in agreement between the left and right body half for the step width and step length is reduced. Abbreviations: RPC: reproducibility coefficient (1.96*SD); CV: coefficient of variation (SD in % of the mean value); ICC(A,1): intraclass correlation coefficient for absolute agreement. Subject averages across steps (AV) and sides (AVG) are reported for comparison with previous publications. Measures in bold can be compared to previously reported systems [1, 2].

| Parameter | Our System | VICON | Pearson's correlation |  |  | Bland-Altman |  |  | ICC(A,1) |
| --- | --- | --- | --- | --- | --- | --- | --- | --- | --- |
| | Mean ± SD | Mean ± SD | R | P | N | Bias | RPC | CV (%) | |
| Stride length L (cm) | 146.08 ± 8.65 | 145.36 ± 8.49 | 0.970 | <0.001 | 267 | 0.71 | 4.16 | 1.46 | 0.966 |
| Stride length R (cm) | 146.53 ± 9.47 | 147.04 ± 9.01 | 0.973 | <0.001 | 187 | -0.52 | 4.32 | 1.50 | 0.970 |
| Step length L (cm) | 74.36 ± 5.16 | 73.95 ± 5.36 | 0.833 | <0.001 | 267 | 0.41 | 5.97 | 4.11 | 0.830 |
| Step length R (cm) | 72.01 ± 5.35 | 71.67 ± 4.48 | 0.817 | <0.001 | 281 | 0.34 | 6.05 | 4.30 | 0.803 |
| Step width L (cm) | 9.94 ± 3.88 | 11.51 ± 3.15 | 0.757 | <0.001 | 267 | -1.57 | 4.98 | 23.70 | 0.675 |
| Step width R (cm) | 14.15 ± 3.47 | 11.91 ± 2.97 | 0.817 | <0.001 | 281 | 2.24 | 3.94 | 15.42 | 0.650 |
| AV stride length L (cm) | 146.37 ± 7.93 | 145.66 ± 7.98 | 0.999 | <0.001 | 10 | 0.71 | 0.71 | 0.25 | 0.995 |
| AV stride length R (cm) | 146.43 ± 8.66 | 146.95 ± 8.43 | 0.999 | <0.001 | 10 | -0.53 | 0.75 | 0.26 | 0.997 |
| **AV stride length AVG (cm)** | **146.40 ± 8.23** | **146.20 ± 8.17** | **0.999** | **<0.001** | **10** | **0.20** | **0.55** | **0.19** | **0.999** |
| AV step length L (cm) | 74.48 ± 4.11 | 74.10 ± 5.13 | 0.967 | <0.001 | 10 | 0.38 | 3.05 | 2.09 | 0.946 |
| AV step length R (cm) | 71.99 ± 4.39 | 71.67 ± 3.60 | 0.927 | <0.001 | 10 | 0.32 | 3.36 | 2.38 | 0.914 |
| **AV step length AVG (cm)** | **73.20 ± 4.01** | **72.82 ± 3.96** | **0.999** | **<0.001** | **10** | **0.38** | **0.35** | **0.24** | **0.995** |
| AV step width L (cm) | 9.97 ± 2.64 | 11.55 ± 1.85 | 0.842 | 0.002 | 10 | -1.58 | 2.88 | 13.67 | 0.649 |
| AV step width R (cm) | 14.16 ± 2.18 | 11.91 ± 1.74 | 0.845 | 0.002 | 10 | 2.24 | 2.29 | 8.98 | 0.505 |
| **AV step width AVG (cm)** | **12.13 ± 2.08** | **11.73 ± 1.76** | **0.936** | **<0.001** | **10** | **0.39** | **1.48** | **6.35** | **0.910** |

## Discussion

We have described an improved motion capturing system based on multiple Kinect v2 sensors in the context of gait analysis. The importance of spatial and temporal calibration has been emphasized. Further, we have demonstrated that the human pose estimation algorithm of the Kinect sensor depends on the view angle and how self-occlusions might lead to biased joint position estimates. The presented gait analysis system has successfully been used to record ten healthy subjects while walking. Recordings have concurrently been performed with a VICON motion capturing system for quantitative comparison. Gait parameters have been extracted from both recordings independently. Agreement with the VICON system has been statistically examined for one-sided and two-sided Kinect tracking, revealing much better agreement for two-sided Kinect tracking. Using two-sided tracking we also reach better agreements in step width with the gold standard than previously reported Kinect systems [1].

### Kinect skeleton tracking is sensitive to view angle

To our knowledge, the most detailed report concerning the accuracy of joint position estimation using the Kinect v2 sensor in comparison to another motion capturing system is provided by Wang et al [4]. In addition to comparing different body poses (sitting and standing), they also examined the influence of the view angle (0°, 30° and 60°) and showed that the joint positions of the turned-away body half are less accurate. They pointed out that the likely cause for the decreased accuracy is the increasing occlusion of one body half by the other (self-occlusion) with increasing view angle. In order to examine the influence of self-occlusions and view angle dependence in a single recording, we recorded a person during the double support phase with our gait analysis system using two sensors and opposite view angles (Figure 5). Based on this recording we reproduced Wang et al’s finding and, further, demonstrated that the joint positions are biased towards the surface area that is visible to the respective sensor (Supplementary Material). Hence, we verified Wang et al’s hypothesis that the decreased accuracy is caused by self-occlusions [4]. As consequence, the extracted gait parameters based on one-sided tracking might not only be less accurate but biased depending on the view angle of the sensors.

### Two-sided Kinect tracking improves overall accuracy

During the gait cycle the left leg partially or completely occludes the right leg every now and then when placing the sensors only on the left side. Depending on the view angle of the sensors this might happen exactly when the two feet have maximum distance (double support phase). Since this event is easily identifiable in the time series of the foot positions, this is exactly the phase of the gait cycle which is commonly used to determine the step length and width as well as the stride length. Inaccuracies in the joint position estimates therefore propagate directly into the respective gait parameters. This is well captured by the agreement statistics of the one-sided Kinect setup with VICON. Specifically, for the step length and step width on the right side the agreement with the VICON system is much lower than for those on the left side which are not affected by occlusions (see Table 2). The stride length is affected by occlusions on any side, because every stride consists of subsequent left and right steps. Gold standard motion capturing systems, like VICON, commonly track the person from all sides, minimizing the amount of self-occlusions due to diverse view angles and multiplicity of sensors. In our two-sided Kinect setup, we increased the number of view angles and thereby decreased the possibility of self-occlusions in comparison to one-sided setups. The effect is prominent when comparing the agreement statistics of the two setups. Both analyses are based on the very same recordings and differ only in the sensors that have been used for the reconstruction of the skeleton. Consequently, the agreement with the VICON system is for the two-sided setup better than for the one-sided setup (compare Table 2 and Table 3). Additionally, we do not observe the strong imbalance of agreements for steps on the left and right side anymore. Unfortunately, Geerse et al. [1] and Mentiplay et al. [2] did not report separate agreement statistics for the left and right steps. In most parameters (subjects’ average step length and stride length, averaged across feet, walking speed) our system reaches similar agreement with the gold standard as other Kinect-based systems [1, 2]. However, especially for the step width and time our system provides better results.

### Precise temporal synchronization is essential for overall accuracy

Previously reported gait analysis systems based on multiple Kinect sensors are often quite unspecific with respect to the technical details of the temporal synchronization of the sensors. Since the Kinect v2 sensor requires a separate USB 3.0 controller and the current version of the Kinect SDK (v2.0) does not support multiple Kinect sensors, most setups use dedicated computers, one for each sensor, just like we do. However, it is known that internal clocks of computers do not run in synchrony and even once synchronized the clocks diverge rather quickly. For this purpose, time synchronization protocols have been developed. Probably the best known one is the network time protocol (NTP) which is for example used by the Windows time service to synchronize the local clock of a computer with a remote clock, also known as time server. PTP in contrast is used to synchronize computers in local networks with high precision, for example in distributed control scenarios. Using commercial software that implements PTP, we have ensured steady synchrony between all involved computers before and during the recordings. One might imagine, that even slight phase shifts due to an asynchrony of the clients lead to inaccuracies when averaging across the data received from different clients. Specifically, since gait is to a large extent described by a periodic function, averaging over different phase shifts leads to smoothing in time and space and, ultimately, to reduced temporal resolution. The gait parameters (step time, length and width as well as the stride length) are consequently less accurate in comparison to a gold standard system which ensures high temporal resolution. We believe that the careful way to calibrate our system not only in space but also in time contributes to the high agreement of our system with the VICON system.

### Scalability of the system

The presented system can easily be scaled up using a larger number of Kinect sensors. To our knowledge, so far only systems with up to four Kinect sensors have been published [1, 14] (also see [18]). In contrast, we have shown that our system is able to utilize six Kinect sensors for tracking people during walking. Since we do not transmit any data in real-time while recording, the network traffic is limited to maintaining the TCP/IP connections between the server and clients and synchronizing the computers. Even though there are currently no obvious limitations in scalability, the maximum number of clients needs to be evaluated.

### Conclusions

We have shown that multiple Kinect v2 sensors can be used for accurate analysis of human gait. Important is the spatiotemporal calibration of the system as well as the sensor placement. Tracking from both sides leads to more accurate and less biased gait parameters which leads to excellent agreement with the VICON motion capturing system for the gait parameters step length, step width, step time, stride length and walking speed. The presented spatial calibration algorithm allows flexible sensor placement which makes fast and easy setup in diverse scenarios possible. We have shown that our system in combination with a two-sided setup of sensors provides better statistical agreement with a gold standard MoCap system than previously reported systems [1, 2]. Besides the application for clinical and scientific gait analysis, the presented motion tracking system might also be useful for the realization of virtual reality setups in combination with head-mounted displays such as the Oculus Rift (Oculus VR, LLC.) or HTC Vive (HTC Corporation).

## Acknowledgements

NL would like to thank the German National Academic Foundation for granting a doctoral student fellowship. Financial support for this study has been received from the European Union Seventh Framework Programme (Koroibot FP7-611909, DFG GI 305/4-1, DFG GZ: KA 1258/15-1, CogIMon H2020 ICT-644727) and the Human Frontiers Science Program (RGP0036/2016). The authors declare no competing financial interests.

## Author Contributions

Software & system development: NL BM. Conceived and designed the experiment: NL BM WI. Performed the experiment: BM. Analysed the data: BM NL. Contributed materials/equipment: MAG. Wrote the original draft: NL BM. Reviewed and Editing: BM NL WI MAG. Acquisition of funding: MAG.

